# Phage mobility is a core determinant of phage-bacteria coexistence in biofilms

**DOI:** 10.1101/086462

**Authors:** Matthew Simmons, Knut Drescher, Carey D. Nadell, Vanni Bucci

## Abstract

Many bacteria are adapted for attaching to surfaces and for building complex communities, termed biofilms. The biofilm mode of life is predominant in bacterial ecology. So, too, is exposure of bacteria to ubiquitous viral pathogens, termed bacteriophages. Although biofilmphage encounters are likely to be very common in nature, little is known about how phages might interact with biofilm-dwelling bacteria. It is also unclear how the ecological dynamics of phages and their hosts depend on the biological and physical properties of the biofilm environment. To make headway in this area, here we develop the first biofilm simulation framework that captures key mechanistic features of biofilm growth and phage infection. Using these simulations, we find that the equilibrium state of interaction between biofilms and phages is governed largely by nutrient availability to biofilms, infection likelihood per host encounter, and the ability of phages to diffuse through biofilm populations. Interactions between the biofilm matrix and phage particles are thus likely to be of fundamental importance, controlling the extent to which bacteria and phages can coexist in natural contexts. Our results open avenues to new questions of host-parasite coevolution and horizontal gene transfer in spatially structured biofilm contexts.

## Introduction

Bacteriophages, the viral parasites of bacteria, are predominant agents of bacterial death and horizontal gene transfer in nature (Suttle 2007, Thomas and Nielsen 2005). Their ecological importance and relative ease of culture in the lab have made bacteria and their phages a centerpiece of classical and recent studies of molecular genetics (Cairns et al 2007, Labrie et al 2010, Salmond and Fineran 2015, Samson et al 2013, Susskind and Botstein 1978) and host-parasite interaction (Bohannan and Lenski 2000, Brockhurst et al 2005, Chao et al 1977, Forde et al 2004, Gomez and Buckling 2013, Gómez and Buckling 2011, Kerr et al 2006, Koskella and Brockhurst 2014, Lenski and Levin 1985, Levin et al 1977, Vos et al 2009). This is a venerable literature with many landmark discoveries, most of which have focused on liquid culture conditions. In addition to living in the planktonic phase, many microbes are adapted for interacting with surfaces, attaching to them, and forming multicellular communities (Meyer et al 2012, Nadell et al 2016, O’Toole and Wong 2016, Persat et al 2015, Teschler et al 2015, van Vliet and Ackermann 2015, Weitz et al 2005). These communities, termed biofilms, are characteristically embedded in an extracellular matrix of proteins, DNA, and sugar polymers that play a large role in how the community interacts with the surrounding environment (Dragoš and Kovács 2017, Flemming and Wingender 2010).

Since growth in biofilms and exposure to phages are common features of bacterial life, we can expect biofilm-phage encounters to be fundamental to microbial natural history (Abedon 2008, Abedon 2012, Díaz-Muñoz and Koskella 2014, Koskella et al 2011, Koskella 2013, Nanda et al 2015). Furthermore, using phages to kill unwanted bacteria – which was eclipsed in 1940 by the advent of antibiotics in Western medicine – has experienced a revival in recent years as an alternative antimicrobial strategy (Azeredo and Sutherland 2008, Chan et al 2013, Levin and Bull 2004, Melo et al 2014, Pires et al 2011, Sillankorva et al 2010). Understanding biofilm-phage interactions is thus an important new direction for molecular, ecological, and applied microbiology. Existing work suggests that phage particles may be trapped in the extracellular matrix of biofilms (Briandet et al 2008, Doolittle et al 1996, Lacroix-Gueu et al 2005); other studies have used macroscopic staining assays to measure changes in biofilm size before and after phage exposure, with results ranging from biofilm death, to no effect, to biofilm augmentation (reviewed by (Chan and Abedon 2015)). There is currently only a very limited understanding of the mechanisms responsible for this observed variation in outcome, and there has been no systematic exploration of how phage infections spread within living biofilms on the length scales of bacterial cells.

Biofilms, even when derived from a single clone, are heterogeneous in space and time (Ackermann 2015, Stewart and Franklin 2008). The extracellular matrix can immobilize a large fraction of cells, constraining their movement and the mass transport of soluble nutrients and wastes (Flemming and Wingender 2010, Teschler et al 2015). Population spatial structure, in turn, has a fundamental impact on intra- and inter-specific interaction patterns (Durrett and Levin 1994, Kovács 2014, Nadell et al 2016). Theory predicts qualitative changes in population dynamics when host-parasite contact rate is not a simple linear function of host and parasite abundance (Liu et al 1986), which is almost certainly the case for phages and biofilm-dwelling bacteria under spatial constraint. It is thus very likely that the interaction of bacteria and phages will be altered in biofilms relative to mixed or stationary liquid environments. Available literature supports the possibility of altered phage population dynamics in biofilms (Ashby et al 2014, Gómez and Buckling 2011, Heilmann et al 2012, Scanlan and Buckling 2012, Vos et al 2009), but the underlying details of the phage-bacterial interactions have been difficult to access experimentally or theoretically. Spatial simulations that capture core mechanistic features of biofilms are a promising avenue to begin tackling this problem. Here, we use a simulation approach to study how the biofilm environment can influence micrometer-scale population dynamics of bacteria and phages, highlighting connections between this research area and classical findings from spatial disease ecology.

Existing biofilm simulation frameworks are flexible and have excellent experimental support (Bucci et al 2011, Estrela et al 2012, Estrela and Brown 2013, Hellweger and Bucci 2009, Hellweger et al 2016, Lardon et al 2011, Nadell et al 2016, Naylor et al 2017), but they become impractical when applied to the problem of phage infection. We therefore developed a new simulation framework to study phage-biofilm interactions. Using this approach, we find that nutrient availability and phage infection rates are critical control parameters of phage spread; furthermore, modest changes in the diffusivity of phages within biofilms can cause qualitative shifts toward stable or unstable coexistence of phages and biofilm-dwelling bacteria. The latter result implies a central role for the biofilm extracellular matrix in phage ecology.

## Methods

When phages are implemented as discrete individuals, millions of independent agents can be active in a single simulation space on the order of several hundred bacterial cell lengths. Moreover, the time scale for calculating bacterial growth can be an order of magnitude larger than the appropriate time scale for phage replication and diffusion. These problems create unmanageable computational load for tracking large population sizes when bacteria and phages are modeled in continuous space, as is the case for contemporary biofilm simulations, which are not designed to accommodate these obstacles (Lardon et al 2011). We therefore developed a new framework customized for studying biofilm-phage interactions. To solve these issues, we reduced the amount of spatial detail with which cells are implemented, using a grid-based approach for bacterial biomass calculation. Within each grid node bacteria are considered well-mixed, and their biomass is converted to bacterial cell counts for infection calculations. We also estimate phage Brownian motion by calculating the analytical solution of the diffusion equation and using it as a distribution of the likelihood of finding each phage at each location, thus eliminating the need for calculating each phage’s movement separately. Our model combines (*i*) a numerical solution of partial differential equations to determine solute concentrations in space, (*ii*) a cellular automaton method for simulating biofilms containing a user-defined, arbitrary number of bacterial strains with potentially different properties, and (*iii*) an agent-based method for simulating diffusible phages (Figure 1).

**Figure 1.**
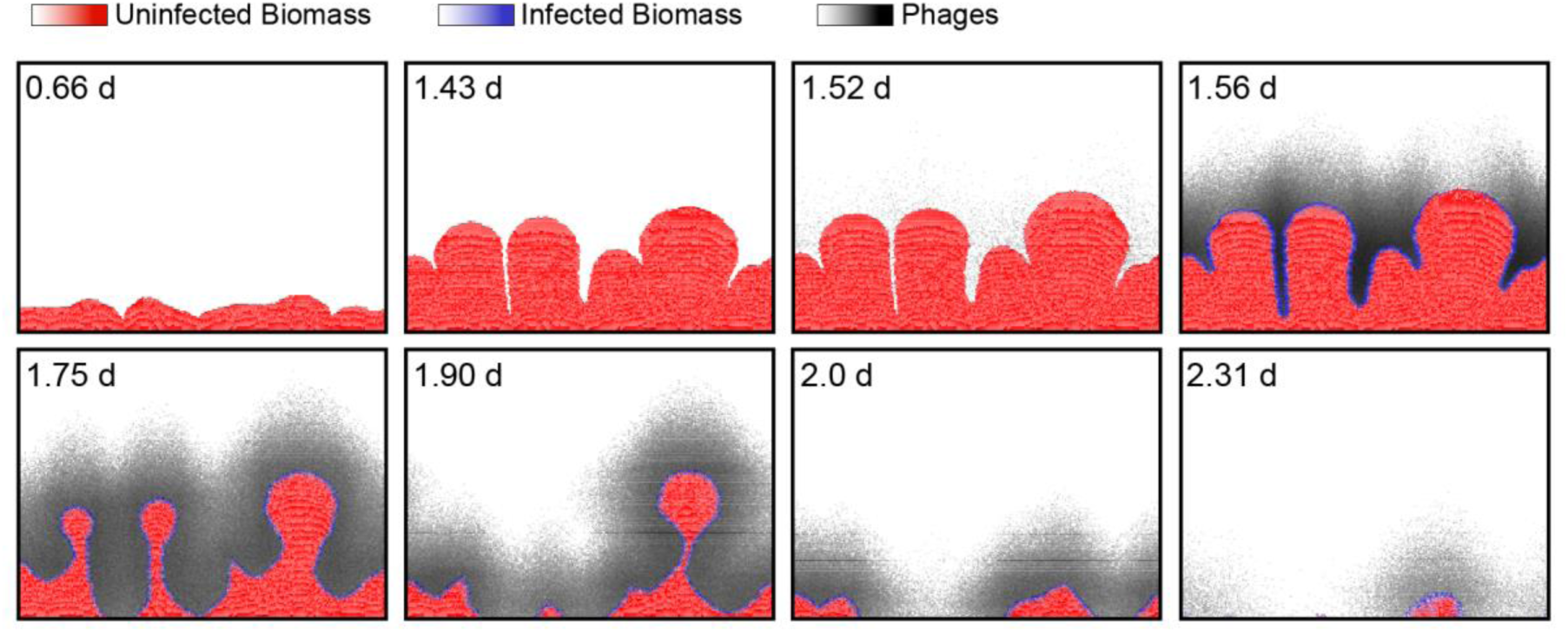
An example time series of simulated biofilm growth and phage infection. For uninfected and infected biomass (red and blue, respectively), the color gradients are scaled to the maximum permissible biomass per grid node (see Supplementary Methods). For phages, the black color gradient is scaled to the maximum phage concentration in this run of the simulation. Any phages that diffuse away from the biofilm into the surrounding liquid are assumed to be advected out of the system in the next iteration cycle. Phages are introduced to the biofilm at 1.5 d. Phage infection proliferates along the biofilm front, causing biomass erosion and, in this example, complete eradication of the biofilm population. The simulation space is 250 μm long along its horizontal dimension.

In each run, the simulation space (250 μm x 250 μm, with lateral periodic boundary conditions) is initiated with cells that are randomly distributed across the basal surface. The following steps are iterated until an exit steady-state criterion is met:

- Compute nutrient concentration profiles
- Compute bacterial biomass dynamics
- Redistribute biomass according to cellular automaton rules (i.e., cell shoving)
- Evaluate host cell lysis and phage propagation
- Simulate phage diffusion to determine new distribution of phage particles
- Assessment of match to exit criteria:
  **Coexistence**: simulations reach a pre-defined end time with both bacteria and phages still present (these cases are re-assessed for long-term stability);
  **Biofilm death**: the bacterial population declines to zero; or
  **Phage extinction**: no phages or infected biomass remain in the biofilm.

As in previous biofilm simulation frameworks, bacteria grow and divide according to local nutrient concentrations, which are calculated to account for diffusion from a bulk nutrient supply (above the biofilm, motivated by flow chamber biofilm culture systems) and absorption by bacteria. Specifically, when nutrients are abundant, most cells in the biofilm can grow. When nutrients are scarce, they are depleted by cells on the outermost layers of the biofilm, and bacteria in the interior stop growing. Cells on the exterior can be eroded due to shear (Alpkvist and Klapper 2007, Chambless and Stewart 2007, Drescher et al 2013, Stewart 2012). Implementing biomass removal by shear is critical in allowing us to study the steady states of the system: without shear-induced sloughing, one is restricted to examining transient biofilm states (Bohn et al 2007, Bucci et al 2011). Sloughing is also required for implementing loss of biomass when phage infections destroy biofilms with rough surface fronts (see below). Bacterial growth, decay, and shear are implemented according to experimentally supported precedents in the literature (Bohn et al 2007, Xavier et al 2004, Xavier et al 2005a, Xavier et al 2005b).

Implementing phage infection, propagation, and diffusion is the primary innovation of the simulation framework we developed. To mimic phages that encounter a pre-grown biofilm after departing from a previous infection site, we performed our simulations such that biofilms could grow for a defined period, after which a single pulse of phages was introduced into the system. During this pulse, lytic phages (Abedon 2008) are added to the simulation space all along the biofilm front. For every phage virion located in a grid node containing bacterial biomass, we calculate the probability of adsorption to a host cell, which is a function of the infection rate and the number of susceptible hosts in the grid node. Upon adsorption, the corresponding bacterial biomass is converted from uninfected to an infected state (Figure 1), and, following an incubation period, the host cell lyses and releases progeny phages with a defined burst size. In the primary analysis below, burst size is fixed at an empirically conservative number, but we also explore robustness of the results to variation in burst size in a supplementary analysis (see Results). Phages move within the biofilm and in the liquid medium by Brownian motion; the model analytically solves the diffusion equation of a Dirac delta function at each grid position to build a probability distribution from which to resample the phage locations.

As the pattern of phage diffusion is probably important for how phage-biofilm interactions occur, we devoted particular attention to building flexibility into the framework for this purpose. The diffusivity of phage particles is likely to decrease when they are embedded in biofilm matrix material, but to what extent phage diffusivity changes may vary from one case to another in natural settings. To study how phage movement inside biofilms influences phage infection dynamics, we introduce a parameter, *Z*_*p*_, which we term phage impedance. For *Z*_*p*_ = 1, phage diffusivity is the same inside and outside of biofilms. As the value of *Z*_*p*_ is increased, phage diffusive movement inside biofilms is decreased relative to normal aqueous solution. Theory predicts that it will be easier for diffusing particles to enter a 3-dimensional mesh maze - which is a reasonable conceptualization of the biofilm matrix - than it is for the same particle to exit the mesh (McCrea and Whipple 1940, Motwani and Raghavan 1995). Our model of phage movement incorporates this predicted property of biofilm matrix material by making it easier for phages to cross from the surrounding liquid to the biofilm mass fraction than *vice versa* (see Supplementary Methods). We also explore the consequences of relaxing this assumption, such that phages can cross from the surrounding liquid to biofilm, and *vice versa*, with equal ease (see Results).

All model parameters, where possible, were set according to precedent in the experimental literature and biofilm simulation literature. There is no experimental system for which all parameters in the framework have been measured, but the key biological parameters used here are, where possible, identical to experimentally measured values for *Escherichia coli* and the lytic phage T7 (Supplementary Table 1). Other key parameters were varied systematically to test for their effects on simulation outcomes (see Results).

To assess the core structure of our simulations, we compared the predictions obtained from a non-spatial version of the framework (i.e., using homogeneous nutrient, bacterial, and phage distributions) with results obtained from an ODE model incorporating the same processes and parameters as the simulations. These trials confirmed that the core population dynamics of the simulations perform according to expectation without spatial structure (see Supplementary Methods and Supplementary Figure S1). A detailed description of the simulation framework and explanation of its assumptions are provided in the Supplementary Methods. The framework code can be obtained from the Zenodo repository: https://zenodo.org/record/268903#.WJho3bYrJHc.

## Computation

Our hybrid framework was written in the Python programming language, drawing from numerical methods developed in the literature (Bell et al 2011, Bresenham 1965, Dijkstra 1959). All data analysis was performed using the R programming language (see Supplementary Methods). Simulations were performed in parallel on the UMass Green High-Performance Computing Cluster. Each simulation requires 4-8 hours to execute, and more than 200,000 simulations were performed for this study, totaling over 100 CPU-years of run time.

## Results

The primary features distinguishing biofilm populations from planktonic populations are spatial constraint and heterogeneity in the distribution of solutes and cellular physiological state, which includes growth rate, and – we hypothesize – phage infection. Our aim here is to identify how these features qualitatively influence bacteria-phage population dynamics in biofilms. We omit the possibility of co-evolution, i.e., we do not consider the origin and maintenance of phage resistance among bacteria, or mutations that alter phage host-range. This simplification was made in order to focus clearly on the mechanisms and impacts of limited movement (of growth-limiting nutrients, bacteria, and phages) on bacteria-phage interaction. The foundation established in this way will be a starting point for understanding the broader problem of eco-evolutionary interplay between phages and their hosts in biofilms.

We began by exploring the different possible outcomes of phage infection in biofilms as a function of phage infectivity, before moving on to a more systematic study of phage transport, phage infection, and bacterial growth rates.

### (a) Stable states of bacteria and phages in biofilms

Intuitively, the population dynamics of bacteria and lytic phages should depend on the relative strength of bacterial growth and bacterial removal, including erosion and cell death caused by phage infection. We studied the behavior of the simulations by varying the relative magnitude of bacterial growth versus phage proliferation. In this manner, we could observe three broad stable state classes in the bacteria/phage population dynamics (Figure 2). We summarize these classes here before proceeding to a more systematic characterization of the simulation parameter space in the following section.

**Figure 2.**
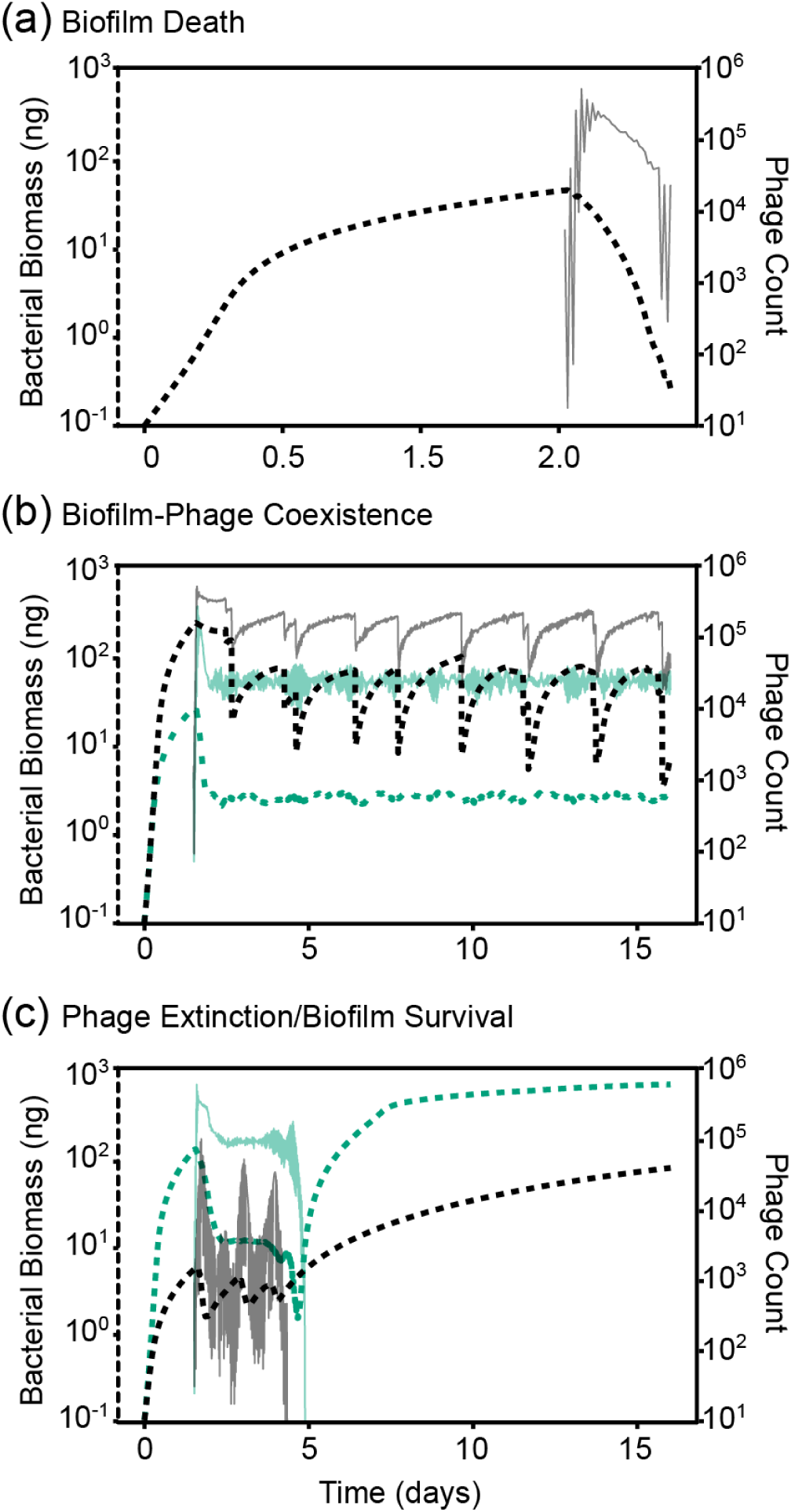
Population dynamics of biofilm-dwelling bacteria and phages for several example cases. For each example simulation, bacterial biomass is plotted in the thick dotted line (left axis), and phage counts are plotted in the thin solid line (right axis) (A) Biofilm death: phages rapidly proliferate and bacterial growth cannot compensate, resulting in clearance of the biofilm population (and halted phage proliferation thereafter). (B) Coexistence of bacteria and phages. We found two broad patterns of coexistence, one in which bacteria and phage populations remained at relative fixed population size (green lines), and one in which bacterial and phage populations oscillated as large biofilms clusters grew, sloughed, and re-grew repeatedly over time (black lines). (C) Phage extinction and biofilm survival. In many cases we found that phage populations extinguished while biofilms were relatively small, allowing the small population of remaining bacteria to grow unobstructed thereafter. Some of these cases involved phage population oscillations of large amplitude (black lines), while others did not (green lines).

#### (i) Biofilm death

If phage infection and proliferation sufficiently out-pace bacterial growth, then the bacterial population eventually declines to zero as it is consumed by phages and erosion (Figure 2A). Phage infections progressed in a relatively homogeneous wave, if host biofilms were flat (Supplementary Video SV1). For biofilms with uneven surface topography, phage infections proceeded tangentially to the biofilm surface and “pinched off” areas of bacterial biomass, which were then sloughed away after losing their connection to the remainder of the biofilm (Supplementary Video SV2). This sloughing process eventually eliminated the bacterial population from the surface.

#### (ii) Coexistence

In some instances, both bacteria and phages remained present for the entire simulation run time. We found that coexistence could occur in different ways, most commonly with rounded biofilm clusters that were maintained by a balance of bacterial growth and death on their periphery (Supplementary Video SV3). When phage infection rate and nutrient availability were high, biofilms entered cycles in which tower structures were pinched off from the rest of the population by phage propagation, and from the remaining biofilm, new tower structures re-grew and were again partially removed by phages (Figure 2B, Supplementary Video SV4). We confirmed the stability of these coexistence outcomes by running simulations for extended periods of time, varying initial conditions and the timing of phage exposure to ensure that host and phage population sizes either approached constant values or entrained in oscillation regimes (see below).

#### (iii) Phage extinction

We observed many cases in which phages either failed to establish a spreading infection, or declined to extinction after briefly propagating in the biofilm (Figure 2C). This occurred when phage infection probability was low, but also, less intuitively, when nutrient availability and thus bacterial growth were very low, irrespective of infection probability. Visual inspection of the simulations showed that when biofilms were sparse and slow-growing, newly released phages were more likely to be swept away into the liquid phase than to encounter new host cells to infect (Supplementary Video SV5). At a conservative maximum bacterial growth rate, biofilms were not able to out-grow a phage infection. However, if bacterial growth was increased beyond this conservative maximum, we found that biofilms could effectively expel phage infections by shedding phages into the liquid phase above them (Supplementary Video SV6). This result, and those described above, heavily depended on the ability of phages to diffuse through the biofilms, to which we turn our attention in the following section.

### (b) Governing parameters of phage spread in biofilms

Many processes can contribute to the balance of bacterial growth and phage propagation in a biofilm. To probe our simulation framework systematically, we used our pilot simulations to choose control parameters with strong influence on the outcome of phage-host population dynamics. We then performed sweeps of parameter space to build up a general picture of how the population dynamics of the biofilm-phage system depends on underlying features of phages, host bacteria, and biofilm spatial structure.

We isolated three key parameters with major effects on how phage infections spread through biofilms. The first of these is environmental nutrient concentration, *N*_max_, an important ecological factor that heavily influences biofilm growth and architecture (Drescher et al 2016, Nadell et al 2010). Importantly, varying *N*_max_ not only changes the overall growth rate but also the emergent biofilm spatial structure. When nutrients are sparse, for example, biofilms grow with tower-like projections and high variance in surface height (Picioreanu et al 1998), whereas when nutrients are abundant, biofilms tend to grow with smooth fronts and low variance in surface height (Nadell et al 2010, Nadell et al 2013, Picioreanu et al 1998). We computationally swept *N*_max_ values to vary biofilm growth from near zero to a conservative maximum allowing for biofilm growth to a height of 250 μm in 24 hours without phage exposure. The second governing parameter is phage infection probability, which we varied from 0.1% to 99% per phage-host encounter. Phage burst size is also important, but above a threshold value (approximately 100 new phages per lysed host), we found that its qualitative influence on our results saturated (Supplementary Figure S2). Lower burst sizes exerted similar effects to lowering the probability of phage infection per host encounter. For simplicity in the rest of the paper, we use a fixed burst size of 100, which is typical for model lytic phages such as T7 (Endy et al 2000).

Our pilot simulations with the framework suggested that a third factor, the relative diffusivity of phages within biofilms, may be fundamental to phage-bacteria population dynamics. We therefore varied phage movement within the biofilm by changing the phage impedance parameter *Z*_*P*_; larger values of *Z*_*P*_ correspond to slower phage diffusivity within biofilms relative to the surrounding liquid.

We performed thousands of simulations in parallel to study the influence of nutrients, infection probability, and phage mobility on population dynamics. In Figure 3, the results are visualized as sweeps of nutrient concentration versus phage infectivity for three values of phage impedance. For each combination of these three parameters, we show the distribution of simulation exit conditions, including biofilm death, phage extinction, or phage-bacteria coexistence. In some cases, biofilms grew to the ceiling of the simulation space, such that the biofilm front could no longer be simulated accurately. To be conservative, the outcome of these cases was designated as “undetermined", but they likely correspond to phage extinction or coexistence.

**Figure 3.**
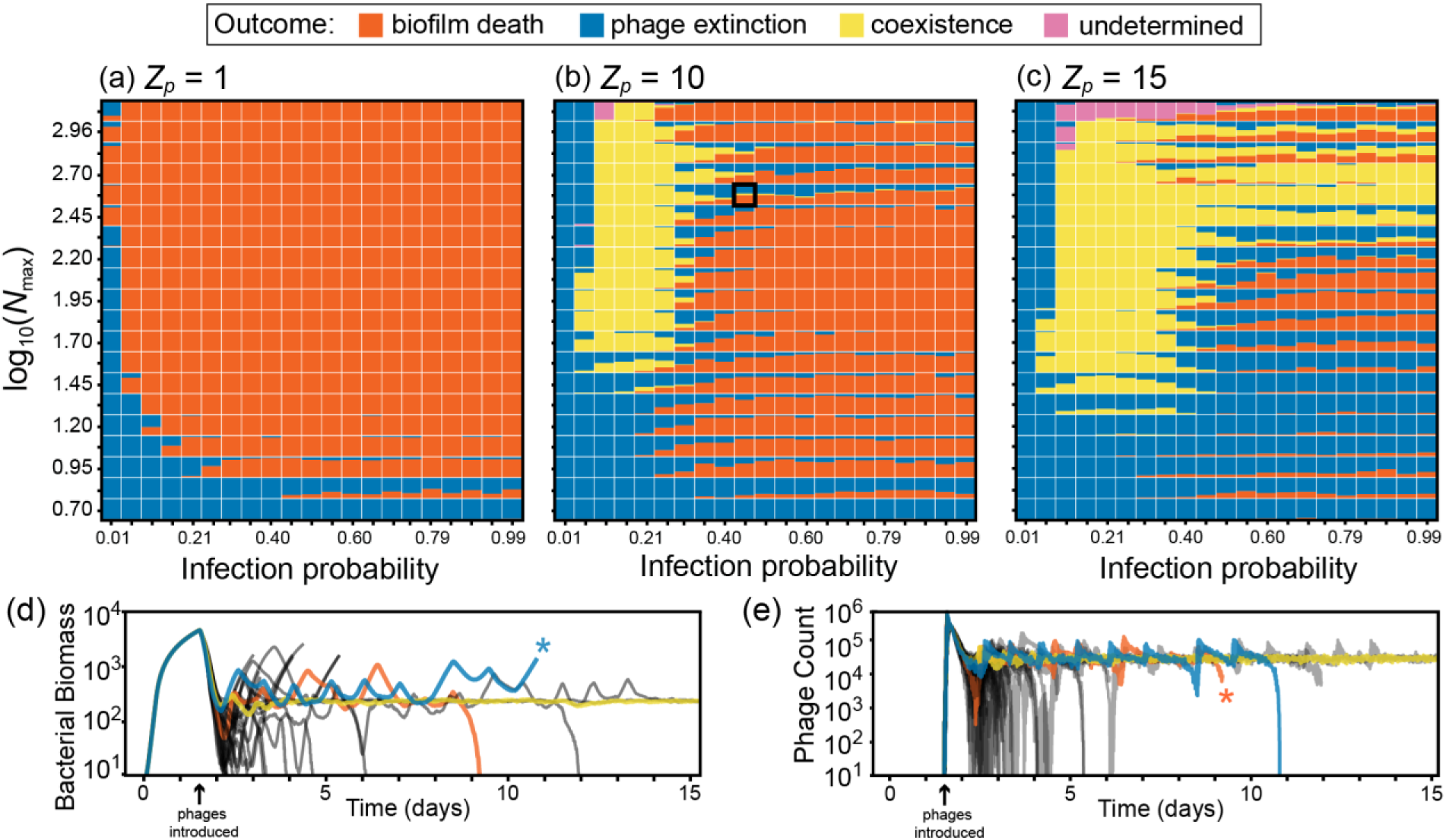
Steady states of biofilm-phage population dynamics as a function of nutrient availability, phage infection rate, and phage impedance. Each point in each heatmap summarizes >30 simulation runs, and shows the distribution of simulation outcomes. Phage extinction (biofilm survival) is denoted by blue, biofilm-phage coexistence is denoted by yellow, and biofilm death is denoted by orange. Each map is a parameter sweep of nutrient availability (∼biofilm growth rate) on the vertical axis, and infection probability per phage-bacterium contact event on the horizontal axis. The sweep was performed for three values of *Zp*, the phage impedance, where phage diffusivity within biofilm biofilms is equivalent to that in liquid for *Zp* = 1 (panel A), and decreases with increasing *Zp* (panels B and C). For *Zp* = [10,15], there are regions of stable coexistence (all-yellow points) and unstable coexistence (bi-and tri-modal points) between phages and bacteria. Traces of (D) bacterial biomass and (E) phage count are provided for one parameter combination at *Zp* = 10 (identified with a black box in panel B) corresponding to unstable phage-bacterial coexistence. We have highlighted one example each of phage extinction (blue), biofilm death (orange), and coexistence (yellow), which in this case is likely transient. In the highlighted traces, asterisks denote that the simulations were stopped because either phages or the bacterial biomass had declined to zero. This was done to increase the overall speed of the parallelized simulation framework. Simulations were designated “undetermined” if biofilms reached the ceiling of the simulation space before any of the other outcomes occurred (see main text).

We first considered the extreme case in which phage diffusion is unaltered inside biofilms (phage impedance value of Z_*p*_ = 1). In these conditions, coexistence does not occur, and bacterial populations do not survive phage exposure unless infection probability is nearly zero, or if nutrient availability is so low that little bacterial growth is possible (Figure 3A). In these latter cases, as we described above, phages either cannot establish an infection at all or are unlikely to encounter new hosts after departing from an infected host after it bursts. Bacterial survival in this regime depends on the spatial structure of biofilm growth, including the assumption that phages which have diffused away from the biofilm surface are advected out of the system. Importantly, using the same experimentally constrained parameters for bacteria and phages, but in a spatially homogenized version of the framework – which approximates a mixed liquid condition – elimination of the bacterial population was the only outcome (Supplementary Figure S1). This comparison further highlights the importance of spatial effects on this system’s population dynamics.

When phage diffusivity is reduced within biofilms relative to the surrounding liquid (phage impedance value of *Z*_*P*_ = 10), biofilm-dwelling bacteria survive infection for a wider range of phage infection probability (Figure 3B). Phages and host bacteria coexist with each other at low to moderate infection probability and high nutrient availability for bacterial growth. Within this region of coexistence, we could find cases where phage and host populations converge to stable fixed equilibria, and others in which bacterial and phage populations enter stable oscillations. The former corresponds to stationary biofilm clusters with a balance of bacterial growth and phage proliferation on their periphery (as in Supplementary Video SV3), while the latter corresponds to cycles of biofilm tower projection growth and sloughing after phage proliferation (as in Supplementary Video SV4). For low nutrient availability, slow-growing biofilms could avoid phage epidemics by providing too few host cells for continuing infection.

As phage diffusivity within biofilms is decreased further (Figure 3C), coexistence occurs for a broader range of nutrient and infectivity conditions, and biofilm-dwelling bacteria are more likely to survive phage exposure. Interestingly, for *Z*_*P*_ = 15 there was a substantial expansion of the parameter range in which biofilms survive and phages go extinct. For *Z*_*P*_ = 10 and *Z*_*P*_ = 15, we also found cases of unstable coexistence in which bacteria and phages persisted together transiently, but then either the host or the phage population declined to extinction stochastically over time (Figure 3D-E). Depending on the relative magnitudes of bacterial growth (low vs. high nutrients) and phage infection rates, this unstable coexistence regime was shifted toward biofilm survival or elimination.

Overall, the tendency of the system toward different stable states in parameter space could be shifted by modest changes in any of the key parameters tested. For example, in Figure 3C, at intermediate phage infectivity, low nutrient availability resulted in biofilm survival. Increasing nutrient input leads to biofilm death as biofilms become large enough for phages to take hold and spread through the population. Further increasing nutrient availability leads to a region of predominant coexistence as higher bacterial growth compensates for phage-mediated death. And, finally, increasing nutrient input further still leads to stochastic outcomes of biofilm survival and biofilm death, with the degree of biofilm sloughing and erosion imposing chance effects on whether biofilms survive phage exposure.

The stochasticity inherent to the spatial simulations provides an inherent test of stability to small perturbations. To assess the broader robustness of our results to initial conditions, we repeated the parameter sweeps, but varied the time at which phages were introduced to the system. We found that the outcomes were qualitatively identical when compared with the data described above (Supplementary Figure S3). Our main analysis in Figure 3 also assumes that key biological parameters are held at one fixed value in the bacterial and phage populations within any single simulation. This is a simplification relative to natural systems (Hellweger and Bucci 2009), in which these parameters may vary from one bacterium and phage virion to another. If this simplifying assumption is relaxed, and maximum bacterial growth rate, phage infectivity, phage burst size, and phage latent period are normally distributed in each simulation run, we again observed qualitatively identical results (Supplementary Figure S4).

### (c) Population stable states as a function of phage diffusivity

The findings summarized in Figure 3 suggest that phage diffusivity (reflected by the phage impedance *Z*_*p*_) is a critical parameter controlling population dynamics in biofilms. We assessed this idea systemically by varying phage impedance at high resolution and determining the effects on phage/bacteria stable states spectra. For each value of phage impedance (*Z*_*P*_ = *1 – 18)*, we performed parameter sweeps for the same range of nutrient availability and phage infection probability as described in the previous section, and quantified the fraction of simulations resulting in biofilm death, phage-bacteria coexistence, and phage extinction (Figure 4). With increasing *Z*_*P*_ we found an increase in the fraction of simulations ending in long-term biofilm survival, either through phage extinction or coexistence. We expected the parameter space in which phages eliminate biofilms to contract to nil as phage impedance was increased. However, this was not the case; the stable states distribution, which saturated at approximately *Z*_*P*_ = *15*, always presented a fraction of simulations in which bacteria were eliminated by phages. This result depended to a degree on the symmetry of phage diffusion across the interface of the biofilm and the surrounding liquid. As noted in the Methods section, theory predicts that phages can enter the biofilm matrix mesh more easily than they can exit, and this is the default mode of our simulations (McCrea and Whipple 1940, Motwani and Raghavan 1995). If, on the other hand, phages can diffuse across the biofilm boundary back into the liquid just as easily as they can cross from the liquid to the biofilm, then as *Z*_*P*_ increases, then biofilm death occurs less often, and bacteria-phage coexistence becomes more predominant (Supplementary Figure S5).

**Figure 4.**
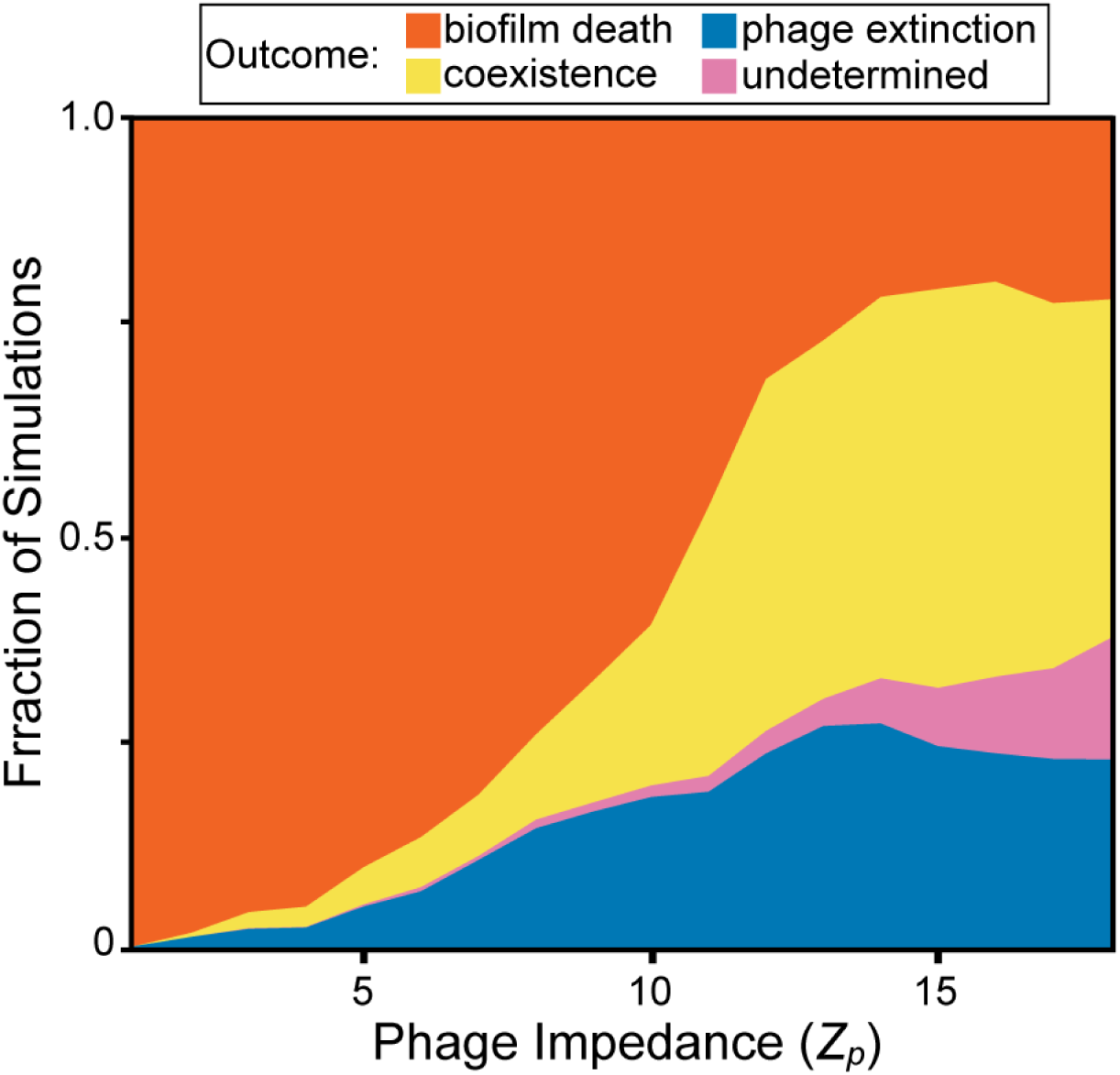
The distribution of biofilm-phage population dynamic steady states as a function of increasing phage mobility impedance within the biofilm. Here we performed sweeps of nutrient and infection probability parameter space for values of phage impedance (*Z*_*P*_) ranging from 1-18. As the phage impedance parameter is increased, phage diffusion within the biofilm becomes slower relative to the surrounding liquid phase. The replication coverage was at least 6 runs for each combination of nutrient concentration, infection probability, and phage impedance, totaling 96,000 simulations. Undetermined simulations are those in which biofilms reached the simulation height maximum before any of the other exit conditions occurred (see main text).

## Discussion

Biofilm-phage interactions are likely to be ubiquitous in the natural environment and, increasingly, phages are drawing attention as the basis for new antibacterial strategies (Abedon 2015). Due to the complexity of the spatial interplay between bacteria and their phages in the biofilm context, simulations and mathematical modeling serve a critical role for identifying and understanding important features of phage-biofilm interactions. Across species and contexts, biofilms are defined by the spatial constraint, altered diffusion environment, and heterogeneous solute distribution conditions created by cells while embedded in an extracellular matrix. Here we developed a new simulation framework that captures these essential processes, and used it to study how they alter the population dynamics of susceptible bacteria and lytic phages.

At the outset of this study, we hypothesized that bacteria might be able to survive phage attack when nutrients are abundant and bacterial growth rate is high. The underlying rationale was that if bacterial growth and biofilm erosion are fast enough relative to phage proliferation, then biofilms could simply shed phage infections from their outer surface into the passing liquid. This result was not obtained, even when nutrient influx and thus bacterial growth were conservatively high. We speculate that for biofilms to shed phage infections in this manner, phage incubation must be long relative to bacterial growth, and/or biofilm erosion must be exceptionally strong, such that biomass on the biofilm exterior is rapidly and continuously lost into the liquid phase. Our results do not eliminate this possibility entirely, but they suggest that this kind of spatial escape from phage infection does not occur under a broad range of conditions.

Biofilms could repel phage attack in our simulations when nutrient availability was low, resulting in slow bacterial growth and widely spaced biofilm clusters. When biofilms are sparse, phage-bacteria encounters are less likely to occur, and thus a higher probability of infection per phage-host contact event is required to establish a phage epidemic. Even if phages do establish an infection, when bacterial growth rates are low, the nearest biofilm cluster may be far enough away from the infected cell group that phages simply are not able to spread from one biofilm cluster to another before being swept away by fluid flow. Note that this observation likely depends on the scale of observation (Levin 1992): in a meta-population context, phage proliferation and subsequent removal into the passing liquid may lead to an epidemic on a larger spatial scale. This caveat aside, our findings are directly analogous to the concept of threshold host density as it applies in wildlife disease ecology (Boots and Sasaki 2002, Holt et al 2003, Keeling 1999, Lloyd-Smith et al 2005, May and Anderson 1979, Maynard-Smith 1974, Rand et al 1995, Satō et al 1994, Webb et al 2007). If host organisms, or clusters of hosts, are not distributed densely enough relative to the production rate and dispersal of a parasite, then epidemics cannot be sustained. Our spatial simulations, which implement the essential biofilm-specific mechanics of bacterial growth and phage infection, can thus recapitulate qualitative features of classical work in spatial epidemiology. This outcome draws concrete links between the microscopic world of phage-host population dynamics and the macroscopic world of disease spread, with results expressed in terms of parameters that are experimentally accessible to microbiologists. We hope that these key concepts may be used in the future as a bridge between researchers studying spatial disease ecology, bacterial biofilms, and bacteriophages.

Our results suggest that coexistence of lytic phages and susceptible host bacteria will occur more readily as phage diffusivity decreases within biofilms, but this outcome also depends strongly on phage infectivity and nutrient flux. In two important modeling studies on phage-bacteria interactions under spatial constraint, Heilmann et al. concluded that coexistence can occur under a broad array of conditions if bacteria are provided with refuges, that is, areas in which phage infectivity is decreased (Heilmann et al 2010, Heilmann et al 2012). An important distinction of our approach is that bacterial refuges against phage infection emerge spontaneously because of the interaction between spatial constraint, biofilm growth, phage proliferation/diffusion, and erosion of bacterial biomass into the surrounding liquid phase. Coexistence of bacteria and phages can be rendered dynamically unstable by modest changes in nutrient availability, phage infectivity, or phage diffusion. In other words, spatial structure is not enough to guarantee phage/bacteria coexistence; rather, given that bacteria and phages are spatially constrained, one must also understand the total balance of biofilm expansion, biofilm erosion, phage infectivity, and phage advection/diffusion in order to understand the system’s population dynamics.

The extracellular matrix is central to the ecology and physiology of biofilms (Branda et al 2005, Dragoš and Kovács 2017, Flemming and Wingender 2010, Flemming et al 2016, Nadell et al 2009, Nadell et al 2015, Nadell et al 2016, Teschler et al 2015). In the simulations explored here, biofilm matrix was modeled implicitly and is assumed to cause changes in phage diffusivity; our results support the intuition that by altering phage mobility and phages’ physical access to new hosts, the biofilm matrix is likely to be important in the ecological interplay of bacteria and their phages (Abedon 2017). A crucial role for the matrix in phage-bacteria interactions is also supported by the common observation that matrix-degrading enzymes are encoded on phage genomes, which indicates that reducing the matrix diffusion barrier is an important fitness currency for phages in natural environments (Chan and Abedon 2015, Pires et al 2016).

Experiments comparing population dynamics of lytic phages and bacteria in well-mixed versus standing liquid cultures indicate that spatial heterogeneity can promote host-parasite coexistence (Brockhurst et al 2006). The biofilm environment shares some conceptual similarity to standing liquid cultures, but is qualitatively different in its details, including sharp gradients of nutrient availability and growth within biofilms, removal of cells from the biofilm system by dispersal, strong diffusion attenuation, and matrix-imposed spatial constraints. Our work lends support to an early suggestion that so-called wall populations on the inner surfaces of culture flasks can promote bacteria-phage coexistence (Schrag and Mittler 1996). The populations described in this work were, almost certainly, biofilms of matrix-embedded cells bound to the flask walls. The details by which this coexistence result occurs have not been clear; there is very little experimental work thus far on the spatial localization and diffusion of phages inside biofilms, but the limited available literature is consistent with the idea that the matrix alters phage movement (Briandet et al 2008, Doolittle et al 1996, Sutherland et al 2004). In biofilms of *E. coli*, the matrix does indeed appear to reduce phage infection (May et al 2011), and recent work with biofilms of *Pseudomonas aeruginosa* grown in artificial sputum further supports the idea that matrix reduces phage susceptibility (Darch et al 2017). Experimental evolution approaches have shown that bacteria and their phages follow different evolutionary trajectories in biofilms versus planktonic culture (Davies et al 2016, Gómez and Buckling 2011, Scanlan and Buckling 2012). Especially compelling in the context of this work, *P. fluorescens* evolves matrix hyper-production in response to consistent phage attack (Scanlan and Buckling 2012). Thinking about phage diffusion and biofilm population structuring will be important not just to the ecological community but also to molecular microbiologists trying to understand the mechanisms underlying phage transport through bacterial populations that are embedded in matrix material.

Here we have identified key properties of phages and their host cells that fundamentally impact population dynamics in bacterial biofilms. To achieve this, some elements of bacteria-phage interaction were not considered here. For instance, we have not implemented co-evolution, though phage and bacterial populations can co-evolve rapidly (Koskella and Brockhurst 2014, Levin and Bull 2004, Perry et al 2015, Thompson 1994, Weitz et al 2005). Selection imposed by phage-mediated killing is responsible for the evolution of diverse host defenses, including altered cell exterior structure, restriction endonucleases, sacrificial autolysis, and the CRISPR-Cas adaptive immune system (Labrie et al 2010). These host defense innovations have, in turn, spurred the evolution of sophisticated host attack strategies on the part of phages (Samson et al 2013). To break ground on the topic of phage-host population dynamics in heterogeneous biofilms, we have set aside the problem of coevolution here; coevolution is undoubtedly important, however, and we expect that biofilm environments will influence it strongly. For example, the typical population sizes of bacteria and phages, as well as their mutual encounter rates, may be dramatically different in biofilms containing tens of thousands of spatially constrained cells, relative to liquid cultures containing tens of billions of well-mixed cells. The time scales and spatial patterns of bacteria-phage coevolution in biofilms may therefore differ substantially from those in liquid culture, which is an important area for future work. We have focused only on lytic phages, but understanding within-biofilm population dynamics of lysogenic phages, which integrate into the genome of infected hosts, often changing their phenotypes and mediating horizontal gene transfer, is also a crucial topic. Overall, we envision that studying bacteria-phage interactions under the unique constraints of biofilm environments will yield important extensions on many fronts of this classical area of microbial ecology.

## Author Contributions

CDN and VB conceived the project; MS, VB, and CDN designed simulations; MS wrote and performed simulations; MS, CDN, VB, and KD analyzed data; CDN, VB, MS, and KD wrote the paper.

## Acknowledgements

We are grateful to Ann Tate and Steve Abedon for comments on an earlier version of the manuscript. C.D.N. is supported by the Cystic Fibrosis Foundation (STANTO15RO), NIH grant P20-GM113132 to the Dartmouth BioMT COBRE, and a Burke Award from Dartmouth College. V.B. acknowledges support from the National Institute of Allergy and Infectious Disease (grant R15-AI112985-01A), and the National Science Foundation (grant 1458347). K.D. is supported by the Max Planck Society, the Human Frontier Science Program (CDA00084/2015-C), the Behrens Weise Foundation, and the European Research Council (716734).

